# Automated assessment reveals extinction risk of reptiles is widely underestimated across space and phylogeny

**DOI:** 10.1101/2022.01.19.477028

**Authors:** Gabriel Henrique de Oliveira Caetano, David G. Chapple, Richard Grenyer, Tal Raz, Jonathan Rosenblatt, Reid Tingley, Monika Böhm, Shai Meiri, Uri Roll

## Abstract

The Red List of Threatened Species, published by the International Union for Conservation of Nature (IUCN), is a crucial tool for conservation decision making. However, despite substantial effort, numerous species remain unassessed, or have insufficient data available to be assigned a Red List threat category. Moreover, the Red Listing process is subject to various sources of uncertainty and bias. The development of robust automated assessment methods could serve as an efficient and highly useful tool to accelerate the assessment process and offer provisional assessments. Here we aimed to: 1) present a machine learning based automated threat assessment method that can be used on less known species; 2) offer provisional assessments for all reptiles - the only major tetrapod group without a comprehensive Red List assessment; and 3) evaluate potential effects of human decision biases on the outcome of assessments. We use the method presented here to assess 4,369 reptile species that are currently unassessed or classified as Data Deficient by the IUCN. Our models range in accuracy from 88% to 93% for classifying species as threatened/non-threatened, and from 82% to 87% for predicting specific threat categories. Unassessed and Data Deficient reptiles were more likely to be threatened than assessed species, adding to mounting evidence that they should be considered threatened by default. The overall proportion of threatened species greatly increased when we included our provisional assessments. Assessor identities strongly affected prediction outcomes, suggesting that assessor effects need to be carefully considered in extinction risk assessments. Regions and taxa we identified as likely to be more threatened should be given increased attention in new assessments and conservation planning. Lastly, the method we present here can be easily implemented to help bridge the assessment gap on other less known taxa.

## Introduction

The IUCN’s (International Union for Conservation of Nature) Red List of Threatened Species [1] is the most comprehensive assessment of the extinction risk of species worldwide [2]. Since its inception in 1964, the Red List has been instrumental in “generating scientific knowledge, raising awareness among stakeholders, designating priority conservation sites, allocating funding and resources, influencing development of legislation and policy, and guiding targeted conservation action” [3]. For example, the 2004 completion of IUCN’s Global Amphibian Assessment, reported a dire state worldwide [4], and led to the creation of organizations dedicated to amphibian conservation and to increased funding for research and conservation policy focused on amphibians [3]. Additionally, the IUCN’s Red List forms a basis for the designation of priority areas for conservation. For example, the Alliance for Zero Extinction [5] works directly with decision-makers to establish protected areas for threatened species represented by a single population, using Red List data.

The Red List assigns evaluated species to categories based on their distribution, population trends, and specific threats [6]. The categories Least Concern (LC) and Near Threatened (NT) are deemed not threatened, while Vulnerable (VU), Endangered (EN), and Critically Endangered (CR) species are deemed threatened. Other species are assessed as Extinct in the Wild (EW), Extinct (EX), or Data Deficient (DD). DD category is assigned to species for which information is insufficient to assign them any of the above categories. Still, most of global biodiversity remains Not Evaluated (NE) by the Red List. This is predominantly due to the laborious nature of Red List assessments, which are based on voluntary expert participation, usually through multi-participant in-person meetings [6]. Importantly, NE and DD species are generally not prioritized for conservation decision making, although Red List guidelines specifically state that they “should not be treated as if they were not threatened” [6]. Even though DD species have been shown to be comparable to CR ones with respect to their levels of overlap with human impact [7]. These assessment gaps [8,9], led to the use of several automated methods to provisionally assess species [10,11]. These methods employ algorithms including phylogenetic regression models [12–14], structural equation models [15], random forests [16,17], deep-learning [18,19], Bayesian networks [20,21], and even linguistic analysis of Wikipedia pages [22]. Most previous attempts (e.g., [12,16,17]) employed a binary classification of threatened (categories CR, EN, VU) versus non-threatened (LC and NT). Few studies attempted to predict specific categories (e.g. [18,19,23]), which are more useful to decision makers as they enable prioritizing among threatened species.

A challenge that remains unaddressed in automated assessment is human decision bias. Biases are introduced by ambiguities in the interpretation of IUCN guidelines by assessors and reviewers, heterogeneity in assessor expertise levels, and personal agendas [24]. The IUCN tries to decrease reliance on subjective expert opinions [2] even employing automated assistance for generating and verifying assessments [11]. However, expert input (and guidance from the IUCN personnel who lead each workshop) remains an important part of the assessment process. Automated methods that ignore such biases in their training data risk reproducing or even amplifying them in their predictions [25].

Reptiles remain the only tetrapod group without comprehensive IUCN assessment. As of July 2021: ∼28% of 11,570 reptile species remain unassessed and ∼14% of those assessed have been classified as Data Deficient [1] Moreover, many of the reptile assessments are more than ten years old rendering them outdated by the IUCN guidelines [1]. This assessment gap is not random. Smaller species, with narrow distributions, located in the tropics, are less likely to have been assessed [8]. Bland and Böhm [26], and Miles [18], automatically assessed some reptile species. Their models predicted ∼20% of NE and DD species are threatened, a similar proportion to those assessed as such (excluding DD). However, in both studies models were trained and validated using a small set of species with a wealth of morphological, ecological, and life history data (which are rare for DD species). Such exercises might provide important information on the mechanisms underlying threat. However, these data-hungry methods are greatly limited in their utility to predict threat status for most DD and NE species, for which few have such data available (e.g., DD and newly described reptiles, most invertebrate taxa). Ultimately, we need methods that will enable precise automated threat assessments of species, that acknowledge different biases and data gaps.

Here, we use robust machine learning to automatically predict IUCN threat categories to all reptile species globally, as a mean to 1) present a new automated assessment framework and 2) provisionally fill the reptile assessment gap. Our methods rely only on readily-available data (mostly geographic ranges, phylogenetic structure, and body mass) and estimate potential effects of assessor or reviewer identities. We use them to assign provisional threat categories to 4,369 reptile species, of which 3,286 currently unassessed and 1,083 currently classified as DD. We further explore global trends in threat across all reptiles and highlight the effects of our new provisional categories on overall patterns in this class. Lastly, we highlight potential sources of biases and incongruences in the assessment process.

## Results

### General model results

We successfully implemented a novel automated assessment method, using the XGBoost algorithm [27], and provided provisional assessment to 4,369 reptile species that were previously not evaluated or assessed as Data Deficient (Supporting Information). Of these 4,369 species, we assessed 1,161 (27%) as threatened (244 as CR, 467 as EN, and 450 as VU), and 3,208 as non-threatened (3,021 as LC and 187 as NT). This is compared to 21% threatened species in the assessed/training dataset (1,375 of 6,520, Chi-square: 26.947, p-value: <0.001).

The complete model, including spatial and phylogenetic autocorrelation, and assessor/reviewer effects, achieved 90% validated accuracy for the binary threatened/non-threatened classification, and 85% accuracy for predicting specific categories (AUC: 0.83, Table 1 and 2). A model excluding assessor/reviewer effects, but retaining spatial and phylogenetic autocorrelation, and one excluding spatial and phylogenetic autocorrelation but retaining assessor/reviewer effects, achieved similar results (Table 1). The model excluding both autocorrelations and assessor/reviewer effects, and the models including either spatial or phylogenetic autocorrelation, were less accurate (Table 1). However, the model obtained the highest accuracies when excluding threatened species classified under criteria other than B from the training data set (Table 1, details below). We predicted threat categories for DD and NE species using the model that excluded assessor/reviewer effects but retained spatial and phylogenetic data, since we cannot know the identity of assessors who will evaluate currently unassessed species. For analyses regarding potential assessor/reviewer effects, we used the complete model. Detailed accuracy metrics are presented in Table 2. The lowest accuracy across models was in separating NT from LC categories (Table 2).

**Table 1.** Comparison of accuracy metrics of eight automated assessment models for classifying reptile species into IUCN threat categories. The “complete” model includes environmental predictors, body mass, spatial and phylogenetic autocorrelations, and assessor/reviewer effects. The species sampling column indicates which species were used in the training of each model, in regard to their threat category and criteria used by IUCN on their assessment. The ’Binary’ task represents the separation of threatened (CR, EN and VU) from non-threatened categories (NT and LC). The ’Specific’ task represents classification into IUCN threat categories. MEM and PEM represent spatial and phylogenetic autocorrelations, respectively. More detailed metrics are presented in Table 2.

| Model | Task | Species sampling | Predictors | Accuracy | AUC |
| --- | --- | --- | --- | --- | --- |
| Complete | Binary | all | Environmental + body mass + PEM<br>+ MEM + assessor/reviewer effects | 0.904 | 0.833 |
|  | Specific | all | Environmental + body mass + PEM<br>+ MEM + assessor/reviewer effects | 0.852 | 0.812 |
| Environment and body<br>mass | Binary | all | Environmental + body mass | 0.877 | 0.784 |
|  | Specific | all | Environmental + body mass | 0.821 | 0.777 |
| Assessor/reviewer<br>effects | Binary | all | Environmental + body mass +<br>assessor/reviewer effects | 0.890 | 0.805 |
|  | Specific | all | Environmental + body mass +<br>assessor/reviewer effects | 0.835 | 0.802 |
| Spatial | Binary | all | Environmental + body mass +<br>MEM | 0.889 | 0.807 |
|  | Specific | all | Environmental + body mass + MEM | 0.825 | 0.791 |
| Phylogenetic | Binary | all | Environmental + body mass + PEM | 0.884 | 0.800 |
|  | Specific | all | Environmental + body mass + PEM | 0.826 | 0.781 |
| Spatial-phylogenetic | Binary | all | Environmental + body mass + PEM + MEM | 0.900 | 0.828 |
|  | Specific | all | Environmental + body mass + PEM + MEM | 0.837 | 0.801 |
| Complete - Criterion B | Binary | Criterion B + NT, LC | Environmental + body mass + PEM + MEM + assessor/reviewer effects | 0.926 | 0.838 |
|  | Specific | Criterion B + NT, LC | Environmental + body mass + PEM + MEM + assessor/reviewer effects | 0.875 | 0.803 |
| Spatial-phylogenetic - Criterion B | Binary | Criterion B + NT, LC | Environmental + body mass + PEM + MEM | 0.915 | 0.800 |
|  | Specific | Criterion B + NT, LC | Environmental + body mass + PEM + MEM | 0.858 | 0.782 |

**Table 2.**
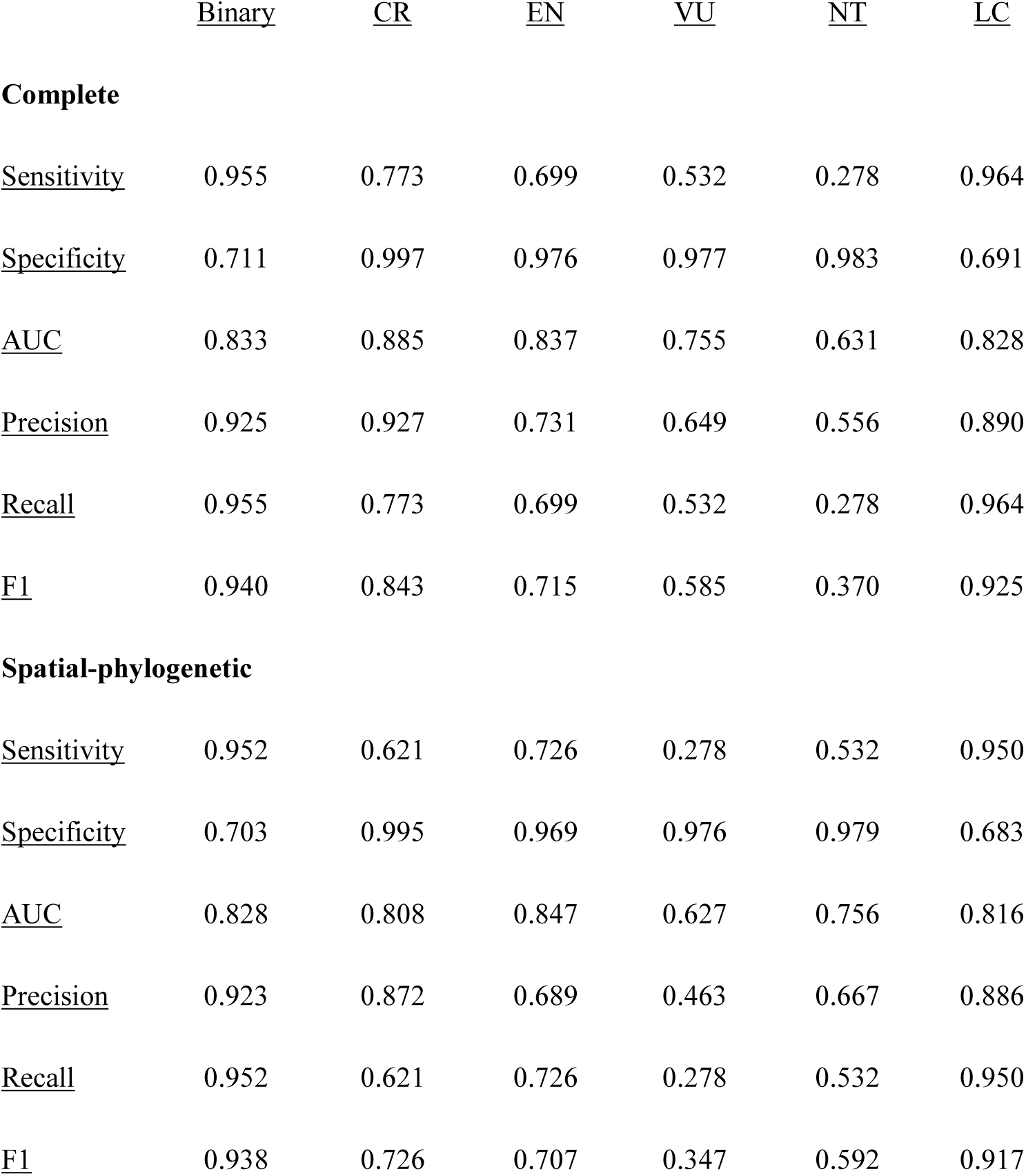
Accuracy metrics of automated assessment models classifying reptile species into IUCN threat categories, under eight different approaches: 1) complete model, accounting for spatial and phylogenetic autocorrelation and assessor/reviewer effects, 2) accounting for spatial and phylogenetic autocorrelation. ’Binary’ represents the separation of threatened (CR, EN and VU) from non-threatened categories (NT and LC). Remaining columns represent the predictive accuracy for assigning species to the five threat categories: CR – Critically Endangered, EN – Endangered, VU – Vulnerable, NT – Near Threatened, LC – Least Concern. See S1 Table for remaining models.

Across different classification tasks and extent of occurrence classes, the average ranking of the importance of feature classes in the complete model was predominantly due to: (1) spatial autocorrelation, (2) assessor effects, (3) phylogenetic autocorrelation, (4) climate. In the model excluding assessor/reviewer effects, the ranking was: (1) spatial autocorrelation, (2) phylogenetic autocorrelation, (3) climate, (4) human encroachment, (for full details on feature importance across models see S2 Table). However, it should be noted that machine learning methods such as XGBoost, are geared primarily towards prediction, not inference [28]. Therefore, we assessed the contribution of features by comparing the predictions of different model configurations (Table 1) and report the feature importance above for their methodological utility. Future studies should explore the underlying mechanisms of reptile extinction risk using more appropriate methods, and clarify which proportion of variation in extinction risk can be attributed to unknown sources of threat correlated in space and phylogeny.

The hyperparameter configuration for the model chosen for predictions is summarized in S3 Table. The features selected for each combination of range size (calculated as extent of occurrence) class and classification task are provided in the Supporting Information. The contribution of each feature class to predictive performance for each combination of range size class and classification task is presented in S1 Fig.

Criterion B for IUCN threat assessments – which is predominantly based on species range sizes [6] – is the most widely used criterion for assigning a threatened status in reptile assessments (74% of species assessed under any criteria). The model including only species assessed as threatened based on criteria B, as well as NT and LC species, was more accurate for both binary (93%, AUC: 0.84, Table 1) and specific categorizations (87%, AUC: 0.80, Table 1). Further excluding assessor/reviewer effects, resulted in similar accuracy (binary classification: 92% accuracy, 0.80 AUC; specific classification: 86% accuracy, 0.78 AUC. Table 1). Despite their higher accuracy, these models tended to misclassify non-criterion B threatened species, assigning them to lower threat categories than observed (S4 Table). This is probably because species are only classified under non-B criteria if such criteria assign them to a similar, or higher, threat category. Thus, we proceeded with models trained on all species for the remaining analysis. Our model correctly classified 93.8% of previously assessed species (6,112 of 6,520 species). The 6.2% misclassified species (408 of 6,520 species) were nearly twice as likely to be assigned to non-threatened categories than to shift in the opposite direction, and generally to shift to less threatened specific categories (S2 Fig). This was consistent in most biogeographical realms, except in the Nearctic and Neotropical realms, in which the numbers were similar for the binary classification (S2 Fig).

### Comparison with previous methods

We compared our method to similar past endeavors. Our method obtained higher accuracy (90%) than methods based on Random Forest (85%) and Neural Networks (79%) (S5 Table). The extreme class imbalance in the dataset greatly hindered both methods, especially Neural Networks (S5 Table), despite the use of supersampling to account for uneven class distributions. In fact, Neural Networks are known to be sensitive to such imbalances [29], while XGBoost is considered more robust to them [27]. While previous methods have incorporated similar predictors to ours and have separately incorporated features such as tolerating missing values, identifying specific IUCN categories, and accounting for spatial and phylogenetic autocorrelation, none does so in combination, as our method does (S6 Table). Our method is also the first to account for assessor bias (Table 3).

### Predictions for Data Deficient and Not Evaluated species

DD and NE species were significantly more likely to be assigned threatened categories than assessed species (DD: 29%, NE: 26%, assessed non-DD: 21% threatened, Fig 1a, S7 Table). DD species were more likely than assessed species to be predicted as VU, EN or CR, and less likely to be predicted as NT or LC. NE species were more likely than assessed species to be VU, and EN, and less likely to be predicted as NT or LC (Fig 1b, S7 Table and S8 Table).

**Figure 1.**
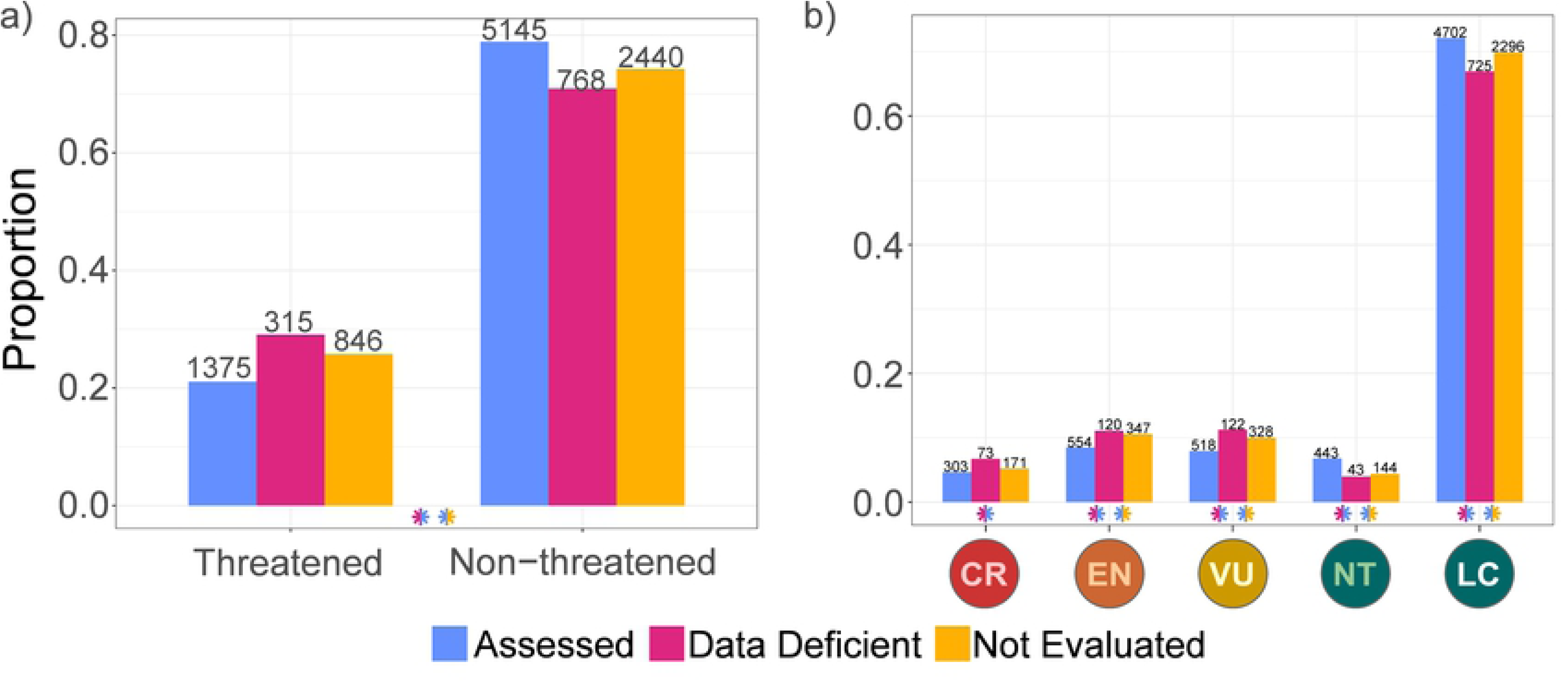
Proportion of reptile species assigned to threat categories by IUCN manual assessment (assessed), and by an automated assessment model (Data Deficient and Not Evaluated). A) Grouping categories into threatened and non-threatened, and B) specific threat categories: CR – Critically Endangered, EN – Endangered, VU – Vulnerable, NT – Near Threatened, LC – Least Concern. Number of species in each category is indicated above each bar. Significant differences in a Pearson’s Χ^2^ test are indicated by asterisks, colored according to which proportions are being compared (S7 Table).

### Phylogenetic and spatial patterns

The proportion of threatened species increased overall for Squamata and Crocodylia but decreased for Testudines (Fig 2, S9 Table), especially in the turtle families Chelidae, Chelydridae, and Kinosternidae. Anguimorph lizards (except Varanidae) threat proportion decreased following our predictions. The three largest lizard clades–Iguania, Scincomorpha, and Gekkota (as well as Lacertoidea except Lacertidae) showed increased threat, as did the largest snake clades (Colubridae, Dipsadinae, Elapidae), and Serpentes (Fig 2, S9 Table). Including predictions for DD and NE species, the proportions of threatened species increased in ecoregions across most of South and North America, Australia, and Madagascar (Fig 3, S10 Table).

**Figure 2.**
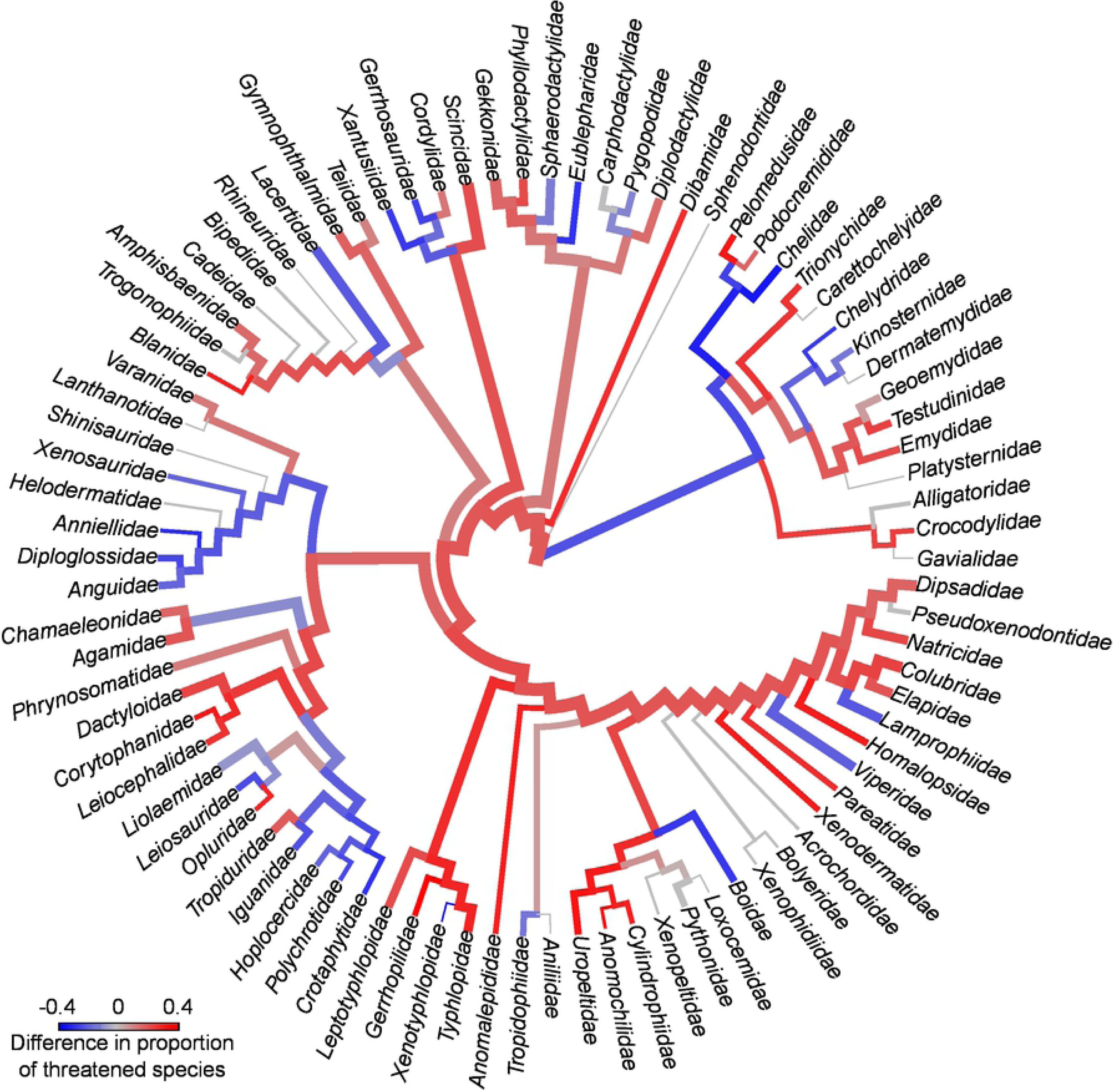
Differences in the percentage of threatened species in reptile families before and after the addition of threat estimates for Data Deficient and Not Evaluated species, obtained from an automated assessment method. Colors in internal nodes represent the difference in percentages for all descendant tips. Trees by Tonini et al. [30] (Squamata) and Colston et al [31] (Archelosauria). The shift between red and blue is proportional to the (symmetric log scale) increase/decrease in threat per branch when using our assessments. Branch widths are proportional to log species richness in each clade. Proportion of threatened species for each family, before and after inclusion of automated assessments are detailed in S9 Table.

**Figure 3.**
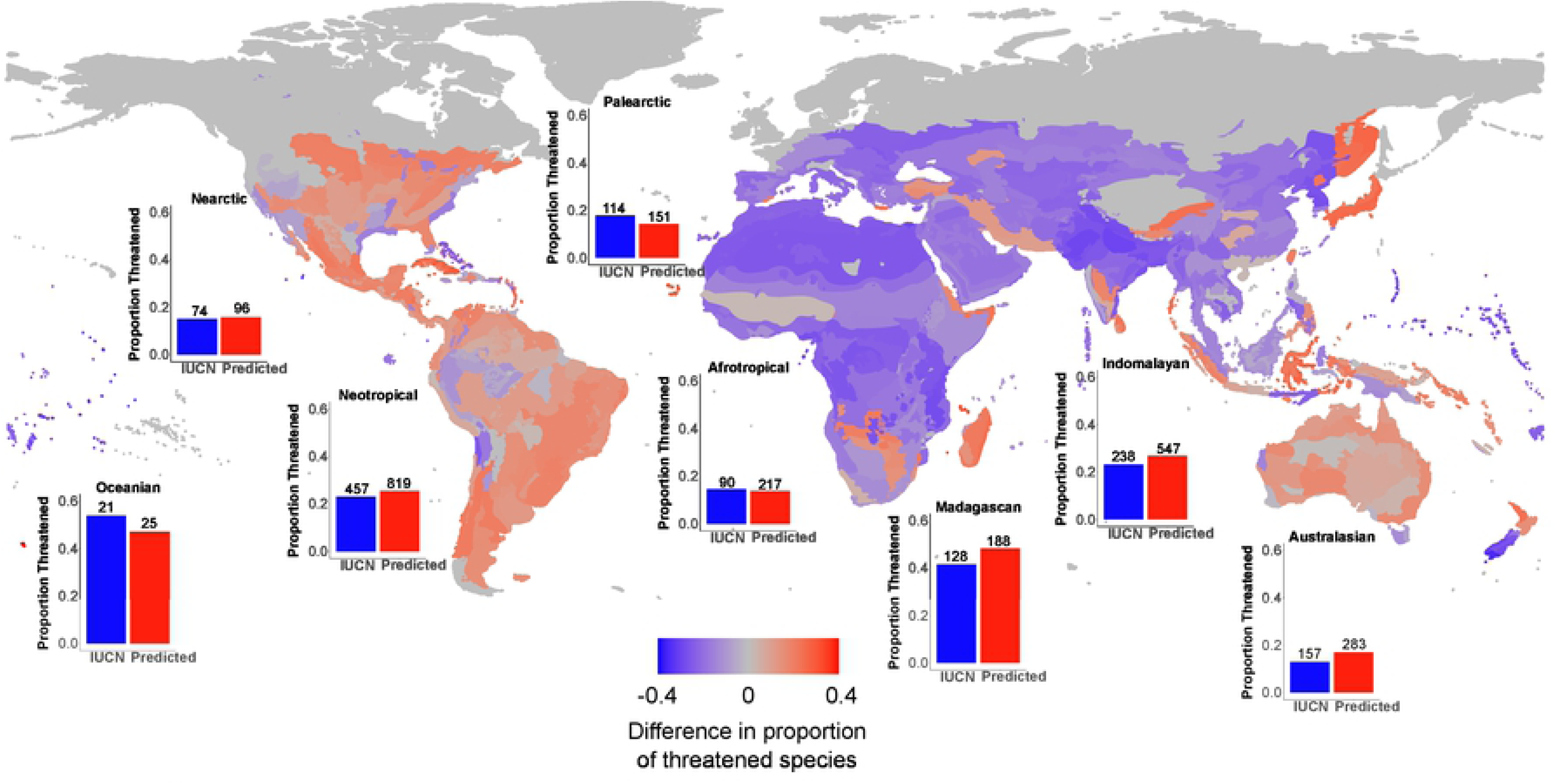
Global spatial changes in the percentage of threatened reptile species resulting from our automated assessments. The spatial data are grouped by WWF terrestrial ecoregions. The shift between red and blue is proportional to the (symmetric log scale) increase/decrease in threat per ecoregion when using our assessments. Bar plots indicate proportion of species in threatened categories for each biogeographical realm, before and after the inclusion of automated assessments.

### Effect of assessor/reviewer identities on predictions

We permuted the identity of assessors and reviewers until we identified the group of assessors and reviewers that would assign each species to the least threatened category possible, while maintaining the other predictors’ values (optimistic scenario) and to the most threatened category possible (pessimistic scenario). Proportions of species predicted as threatened increased from optimistic to observed to pessimistic scenarios for all categories (Fig 4a, S11 Table) and across most biogeographical realms. In the Nearctic and Madagascar, the observed and pessimistic scenarios were similar, and in Oceania no differences were detected (Fig 4b, S12 Table). Species that changed category between the observed assessments and the optimistic scenario move overwhelmingly to a single category (LC), while in the pessimistic scenario, species showed a more diverse distribution of new categories (S3 Fig).

**Figure 4.**
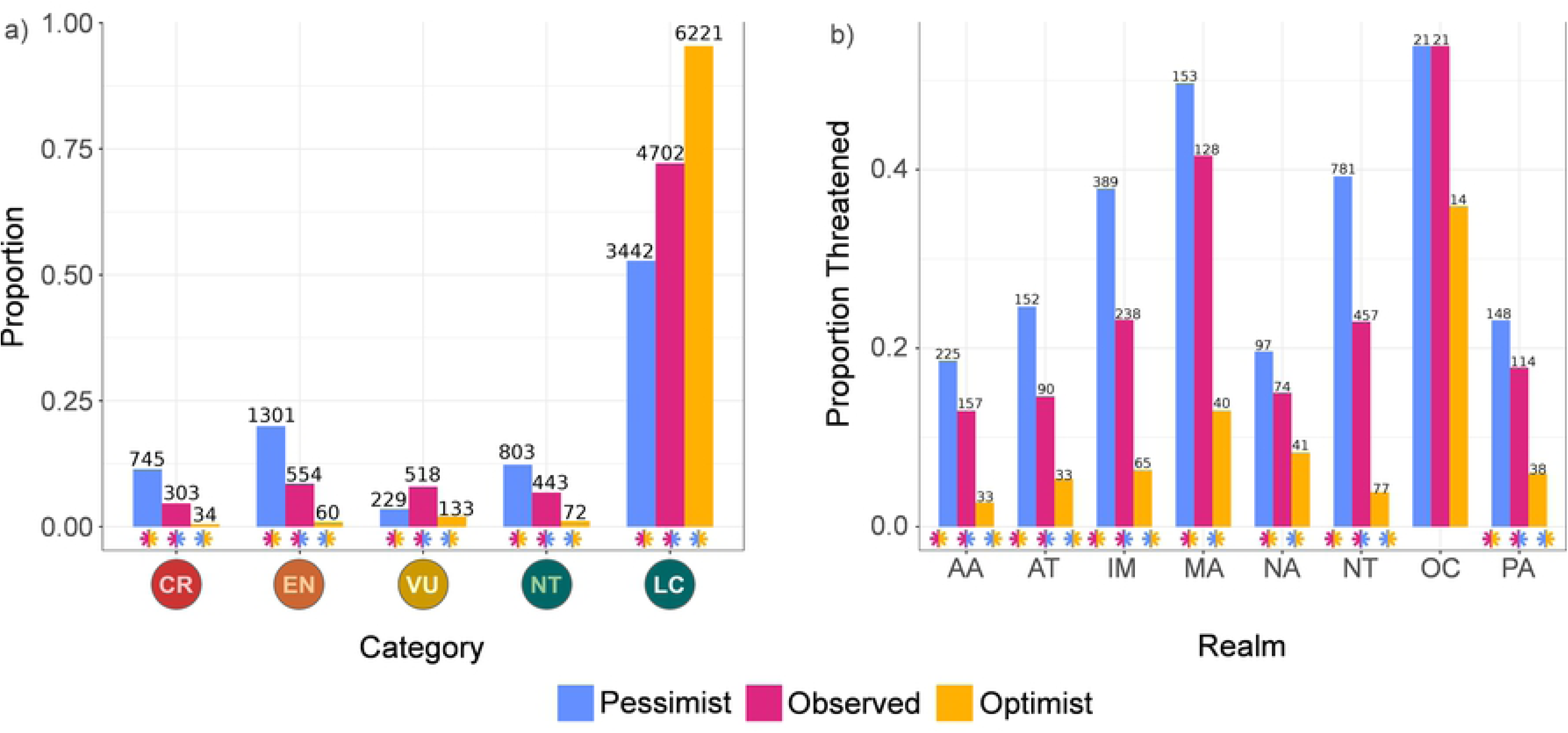
Proportion of threatened reptile species under different assessor bias scenarios. Analysis includes only species that have IUCN assessments (6,520 species) A) Proportion of reptile species assigned to each threat category for the actual IUCN assessments (Observed); proportion expected if the most optimistic group of assessors assessed every species (Optimistic); proportion expected if the most pessimistic group assessed every species (Pessimistic). B) Proportion of threatened species in each biogeographical realm for Observed, Optimistic and Pessimistic assessments. AA – Australasian, AT – Afrotropical, IM – Indomalayan, MA – Madagascan, NA – Nearctic, NT – Neotropical, OC – Oceanian, PA – Palearctic. Significant differences in a Pearson’s Χ^2^ test are indicated by asterisks, colored according to which proportions are being compared (S11 Table).

## Discussion

Our model successfully assigned IUCN categories, and threat statuses, to the 40% of the world’s reptiles that currently lack published assessments or are classified as Data Deficient. Our novel modeling approach enabled classifying specific threat categories with high accuracy using only readily available data (ranges and body sizes). Our methods also gained better accuracy then previously explored methods (S5 Table). We predicted that the prevalence of threatened reptile species is significantly higher than currently depicted by IUCN assessments. This pattern is widespread across space and phylogeny. Our results show that, while high prediction accuracy can be achieved without accounting for assessor/reviewer identities, the identity of assessor/reviewers greatly affects predictions.

### General model results

The classification accuracy of more extreme categories (CR, EN, LC) was higher than categories straddling the threatened/non-threatened threshold (VU and NT, S1 Table). This likely reflects ambiguities inherent to the assessment of borderline cases, while extreme cases are easier to identify. This is compounded in the hardest category to predict (NT), as there are no distinct quantitative thresholds for NT as there are for threatened categories (although guidance is given by the IUCN on how NT should be assessed [6]). Such thresholds are a primary factor for assigning criterion B threat designations (and our modeling). Misclassifications of assessed species tended towards less threatened categories (S2 Fig), indicating that our predictions of unassessed species may actually be more optimistic than the true state of threat for reptiles.

Nine species classified as CR by IUCN were considered LC by our model. Some of these have fragmented ranges (*Spondylurus lineolatus*, *Liolaemus azarai*, *Emoia slevini*), which might have caused the overestimation of their extent of occurrence, since this measure does not account for such range discontinuities. In such cases, species area of occupancy can greatly differ from extent of occurrence, which was the metric used in our models. Thus, species evaluated under area of occupancy criteria might be harder to capture in our model. Small and fragmented ranges can also be more unstable, which might result in discrepancies between the datasets used to train the model. GARD ranges data represents historical ranges, including parts of the range from which populations may have been extirpated. This might cause some of the discrepancies observed. For example, the GARD database includes range fragments of *S. lineolatus* that are classified as possibly extinct in the IUCN database.

Other species classified as less threatened by the model suffer from threats such as invasive species (*Liolaemus paulinae, Cyrtodactylus jarakensis*), quarrying (*Homonota taragui, Cyrtodactylus guakanthanensis*), tourism (*Calamaria ingeri*) and fires (*Bellatorias obiri*) which are not accounted for in our modeling. Although some of the human encroachment features included might act as proxies for such threats, some local stressors will escape this approximation.

Four species (*Tropidophis xanthogaster*, *Cubatyphlops perimychus*, *Celestus marcanoi* and *Chioninia spinalis*) were classified as LC by IUCN, but as CR by our model. All are small ranged species located in protected areas. Protected area effects and local population dynamics may not have been captured by our model in rare cases, leading to occasional overestimation of threat. Alternatively, actual assessments may have been inconsistent with most of the Red List. These are poorly known species, their IUCN assessments read: “*while threats have been identified, these are presently localized*” (*T. xanthogaster)*; *“the limited information available indicates that it is able to adapt at least to certain forms of disturbance”* (*C. perimychus)*; *“there is no information about its population… Further research into its distribution, abundance, and population trends should be carried out to have more knowledge about how the threats are impacting the species”* (*C. marcanoi)*. This lack of information opens room for the introduction of biases, such as overly optimistic assessors overlooking important threats. All four species classified as LC by IUCN and CR by our model have extremely restricted ranges and are endemic to islands with high proportion of threatened species. Thus, we suggest these species may be more threatened than currently depicted in the Red List and would benefit from reassessment. Similar attention should be given to all species that moved to a more threatened category in our assessment (Supporting Information). We recommend a strongly precautionary approach to translating such disparities into conservation action.

Other than differences in range sizes between GARD and IUCN datasets, misclassifications of species as less threatened than assessed by the IUCN may be due to species meeting Red List criteria other than B, as their exclusion led to higher model accuracy. These criteria are mostly based on data on population sizes and trends, which are unavailable for most reptile species. Population dynamics are difficult to approximate using remotely sensed predictors [32], such as the ones used in most automated assessment methods. Excluding species classified as threatened under non-B criteria from model training caused their threat to be severely underestimated (S4 Table). This highlights that the inclusion of population size and trend data in the model can only increase the level of predicted threat compared to the result expected under criterion B only, mimicking the IUCN assessment process.

Nevertheless, most of our modeled classifications (for assessed species) are the same as the IUCN ones (94%, 6,112 of 6,520). The provisional assessments we obtained for unassessed and DD species can therefore be used to accelerate the assessment process, providing a baseline category from which assessors can begin their evaluations. They can then complement it (or change it) with more specific information, when available. The modeled categories can also be useful as guidance when resolving contentious assessments, by providing expected categories based on general threat patterns across taxa or regions. For species that lack any information other than the modeled category, the automated assessments could provide a provisional category under criterion E (Extinction Probability Analysis) or under a newly made criterion (that can be named e.g., E2 or F) for automated assessments, with clear indication of the provisional, modelled, status of the assessment. We recommend that models are associated with expert input especially for species in borderline categories (VU and NT), for which the automated assessment was less reliable. We applied our methods to all DD and NE reptiles globally. In practice, our method can also be applied to regional- and country-level assessments. This is the scale at which national red lists, which support many country-level conservation decisions, are made [33]. Nevertheless, in some regions, challenges, such as lack of resources or standardized methods for regional assessments, are especially salient [34]. Provisional assessments provided by automated methods such as ours can also be used to inform conservation policy and action on DD and NE species, which are currently often given little weight, if any.

### Predictions for Data Deficient and Not Evaluated species

Our results suggest DD species are more likely to be threatened than categorized species, adding to growing evidence in that regard [7,13,16,35–37], but unlike previous automated assessments for reptiles [18,26]. However, it is important to note that previous assessments have drawn on different data sets, both with respect to predictors used and level of extinction risk, as range maps and threat categories have since been updated. We further found that NE reptiles (similar to DD species) are more likely to be threatened than categorized species–supporting the urgency of previous calls for a comprehensive reptile assessment [8]. Our method relies on extent of occurrence maps, which were used as a hierarchical classifier in modeling. Non-DD assessed species have an extent of occurrence that is 16% larger, on average, than DD and NE species (F-value: 6.93, p-value: 0.009). For NE species this may be caused by them being recently described (i.e., later then a workshop on the fauna of the area they inhabit was conducted), and thus having small extent of occurrence. Taxonomic revision resulting in species splits will also give rise to NE species with small extents of occurrence. With such alarmingly high levels of predicted threat, we recommend that decision makers take a cautious stance and assign DD and NE species similar priority as threatened species, unless evidence to the contrary is available (e.g., having been assigned a non-threatened category by an automated assessment).

Data Deficient species may have incomplete distribution records or suffer from taxonomic uncertainties (although only 69 of the 1,083 DD species examined here were classified as such due to taxonomic uncertainty), which might cause their ranges to be underestimated. On the other hand, many truly rare and small-ranged species lack information to be assigned a threat category. It is useful to provide DD species with provisional assessments because they often cannot be used for conservation prioritization [37]. Thus, it is safer to assume that DD species indeed have the ranges from which they are presently known, rather than risking leaving very threatened species in an unprioritizable category [7].

### Phylogenetic and spatial patterns

Our results revealed an overall decrease in the proportion of threatened turtle species after the addition of our predictions for DD and NE species (Fig 2). This could be due to the more complete assessment of turtles than of squamates. Data on population sizes and trends are much more readily available for testudines in than for squamates [38]. Only 19% of squamates were classified as threatened based (at least in part) on criteria other than B – compared to 83% of turtles. The proportion of threatened species tended to increase in squamate groups, especially in small, fossorial, rare and endemic groups (Fig 2, S9 Table), which is consistent with previously reported patterns of data deficiency [8]. Our method is thus better suited for data poor clades than for extremely data-rich ones. The latter have already been assessed or are easy to assess, but the former comprise most of global biodiversity. Thus, our method could be especially useful for other data poor and underassessed groups, such as most invertebrate clades.

Our results suggest that the world’s unknown and rich biodiversity is at even greater risk than previously perceived. This finding adds to accumulating evidence that geographical and phylogenetic patterns of threat and knowledge gaps are mostly congruent [9]. We further found that the proportion of threatened species increases in most ecoregions in the Americas, Australia, and Madagascar but decreases in most of Africa and Eurasia. This could be driven by a taxonomic effect, as many of the families predicted to increase in threat proportion are especially diverse in the Americas, Australia, and Madagascar (e.g., Dactyloidae, Diplodactylidae, Dipsadidae, Elapidae, Phrynosomatidae, Scincidae; Fig 2). Assessments of regions and taxa we identified as likely to be more threatened should be given increased attention in new assessments and conservation planning.

### Effect of assessor/reviewer identities on predictions

Our models achieved high levels of accuracy even without accounting for assessor/reviewer effects (Table 1). Nonetheless, the composition of assessors may greatly influence predictions across all categories (Fig 4a, S3 Fig, S8). A possible explanation for this pattern is that such effects could be implicitly accounted for in spatial and phylogenetic autocorrelation since assessors usually assess only particular taxa and locations (Table 1). For example, if a group of assessors worked mostly on assessment of South American turtles, the biases they introduce might be accounted for the spatial dependency associated with South America and phylogenetic dependency associated with Testudines.

For all realms except Oceania, we found assessor and reviewer identities affected IUCN assessments. The effect of permuting assessor/reviewer identities suggested that observed assessments were similar to those expected if all species were evaluated by the most pessimistic assessors/reviewers in Madagascar, and the Nearctic realms. The lack of effects for Oceania (Fig 4b, S12 Table) is likely due to the small number of species in this realm, and to the few people assessing them. Several recommendations have been made to address assessor bias, including the need for thorough documentation and divulgation of contentious assessments, so they can be used for training and guideline refinement; and training assessors, specifically addressing handling uncertainty and assessor’s attitudes to risk [11,24]. We further recommend that the IUCN, and local or regional agencies wishing to assess extinction risk of species or populations, to: 1) conduct regular automated assessments of previously assessed species, followed by examination of discrepant cases and reassessment if necessary; 2) create a new criterion specifically tailored to provisional automated assessments, as long as the provisional status of the assessment is always clearly indicated; 3) recommend that data scientists are present during the assessment proces**s**, for the production and interpretation of analytical inputs such as automated assessments.

We also recommend, as further research avenues, the development of 1) analytical methods to identify which assessment criteria and subcriteria are more subject to ambiguities, and how can they be refined; 2) applications for quick automated assessments using methods such as the one proposed here; 3) automated assessment methods specifically geared towards modeling population sizes and trends (e.g., based on spatial distribution of threats such as land use changes, climate change, invasive species ranges, and hotspots of wildlife trade), to evaluate species using criteria other than B.

Automated assessments have great potential for reducing bias and uncertainty, by providing a category, based on general patterns in the data, which assessors can use as a starting point to add their specific knowledge to, thus accelerating, standardizing, and refining IUCN’s assessment process. Such improvements would certainly contribute to better informed decision making and increased conservation efficiency. We have shown that accurate predictions can be made without explicitly accounting for assessor/reviewer effects. Previous automated assessments, which reported high levels of accuracy without accounting for assessor/reviewer effects, showed much lower accuracy when their predictions were confronted with manual assessments [26]. Biases from past assessments can be indirectly captured by algorithms and be accurately incorporated in predictions, but biases from future assessments could fall outside the scope of the training data. The contingency of manual assessments on assessor identities makes automated assessments more reliable, but those are also subject to many sources of uncertainty [39,40]. Moreover, since automated methods are trained using previous manual assessments, they risk carrying over the biases of past assessors. Automated methods, that explicitly incorporate uncertainty into their predictions (e.g., [21]), are a promising avenue for future development, and they should explicitly account for assessor/reviewer effects. Overall, automated assessment can be a useful tool for provisional prioritization and assessment acceleration but should be viewed critically.

## Conclusions

We show that, with the inclusion of estimates for DD and NE species, reptiles globally emerge as more threatened than the IUCN Red List currently depicts. This underestimation is widespread across space and phylogeny. Our automated assessments successfully and accurately captured the threat categories and could be widely used for generating provisional assessments for numerous taxa awaiting assessments. We nonetheless recommend that special attention is paid to population declines, which are less well captured by our model and result in it being conservative in assigning threat categories. From a precautionary principle perspective our results also support the notion that DD and NE species should probably be provisionally considered as threatened – if no other sources are available. While IUCN assessments will continue to be the gold-standard for categorizing species threat, we recommend caution is necessary and that assessor/reviewer effects should be considered when using them. Altogether, our models predict that the state of reptile conservation is far worse than currently estimated, and that immediate action is necessary to avoid the disappearance of reptile biodiversity.

## Materials and methods

### Data acquisition

We obtained distribution estimates of 10,889 terrestrial and freshwater reptile species (94% of the 11,570 currently recognized species) from an updated version of the Global Assessment of Reptile Distributions (GARD 1.7 - http://www.gardinitiative.org [41]). We extracted summary values for a suite of parameters obtained using the overlap of each species’ range with five classes of remotely sensed predictors. These include climate (76 features), human encroachment (45 features), biogeography (26 features), topography (9 features), ecosystem productivity (8 features), as well as the latitudinal centroid of each species’ distribution. Predictors and metadata are summarized in the Supporting Information. We added to these predictors species-level data on body mass and insularity assembled from the literature as part of the GARD initiative ([42]; see the Supporting Information). As other biological attributes are harder to come by (and consequently had a lot of missing values for our reptile species) we only included body mass as a species-level biological attribute. We used these data, together with measures of spatial and phylogenetic autocorrelation, and assessor and reviewer effects, to model IUCN threat categories using a recent gradient boosting algorithm (details below). While we used the best available data sources, with the most complete coverage, there might still be geographical biases in their precision. Such biases are likely to occur in any exploration of such a wide scope, and we believe they do not detract from our method. We set aside 20% of species for validation. We used the 15 March 2021 IUCN reptile assessments [1]. All data sets were standardized to the taxonomy of the March 2021 version of the Reptile Database [43], with the input of experts from the GARD initiative. All analysis were conducted in R 4.0.3 [44].

### Incorporating spatial and phylogenetic autocorrelation

We used Moran’s Eigenvector Maps and Phylogenetic Eigenvector Maps to represent spatial and phylogenetic structure in our models [45,46]. The main advantage of these techniques is that they can be incorporated in modern machine learning methods, such as XGBoost [27] (description below). Eigenvector methods have been criticized for requiring the omission of part of the autocorrelation structure and not explicitly incorporating an evolutionary model [12,47]. Some of these critiques have since been resolved [46] and are less relevant in our case as we simply use eigenvectors as proxies for broad scale predictors of threat (see also [48]).

We used the GARD distribution dataset to calculate Moran’s eigenvectors, employing R package ’adespatial’ [49]. We intersected species distribution polygons as neighbors and weighted the neighborhood matrix by inverse centroid distances calculated with function ’nbdists’ from package ’spdep’ [50]. To calculate phylogenetic eigenvectors, we used package ’MPSEM’ [51], and the phylogenies from Tonini et al. [30] for Squamata and Colston et al. [31] for Testudines and Crocodylia. We assumed a Brownian motion model of trait evolution. Species with distribution data but no phylogenetic information (n = 167) were assigned an NA value for all phylogenetic eigenvectors. High positive eigenvalues are associated with autocorrelation at broader scales [45,46]. Since autocorrelation at small scales does not provide information on the entire structure [52], we used eigenvalues to reduce the number of eigenvectors, retaining only eigenvectors with eigenvalues larger than 10% of the eigenvalue of the first eigenvector. This left us with eigenvectors corresponding to autocorrelation structures deeper in the trees and across broader spatial scales. Following this procedure, we retained 236 spatial and 78 phylogenetic eigenvectors.

### Incorporating assessor and reviewer effects

We obtained the identity of 983 assessors and 192 reviewers for all evaluated reptiles on the 15 March 2021 using R package ’rredlist’ [53]. Many of these assessors and reviewers worked together on the assessments of different species in different combinations. To address this, we used an autocorrelative approach similar to our spatial autocorrelation detection/correction method, to incorporate potential assessor/reviewer effects in our models. We considered assessors/reviewers that worked together on a species assessment to be neighbors in the neighborhood matrix, with the number of species each pair assessed together as the weight of each pair’s association. Therefore, frequently associated assessors had more similar scores than those that associated occasionally. Assessors/reviewer scores were averaged for each eigenvector on each species. Therefore, species that were evaluated by a similar set of assessors/reviewers had more similar scores than species evaluated by more distinct sets of assessors/reviewers. We performed *a priori* selection based on eigenvalues, as described above, using the same thresholds, which resulted in 216 eigenvectors being retained for assessors and 39 for reviewers.

### Modeling threat

We used the XGBoost regularizing gradient boosting classification framework in our modeling of threat categories. XGBoost is a recently developed machine learning algorithm that combines computational efficiency, versatility, and high levels of accuracy [27]. It is considered a state-of-the-art machine learning technique and is a popular choice for machine learning competitions [54]. Another advantage of XGBoost is its “Sparsity-aware Split Finding” algorithm which enables effective classification of entries containing missing data [27]. XGBoost is also robust to imbalanced datasets [27], as is the case for reptile threat categories, 72% of which are currently classified as LC [1]. We implemented this algorithm using the R package ’xgboost’ [55] To compare model accuracy and efficiency across algorithms, we further fit a similar model using the AdaBoost algorithm [56], implemented in the R package ’adabag’ [57]. This approach obtained lower accuracy (see Supporting Information).

The range size of a species (as measured by extent of occurrence) can be used as an important *a priori* consideration for the assessment process, since most reptiles are assessed under criterion B. Consequently, we first separated species into the range size classes used in the IUCN Red List B criterion (over 20,000 km^2^, between 20,000 km^2^ and 5,000 km^2^, between 5,000 km^2^ and 100 km^2^, under 100 km^2^). This initial separation enabled different hyperparameter tuning, feature selection and model fitting for each extent of occurrence class. Next, we used a decision tree (Fig 5) involving four hierarchical classification tasks for each extent of occurrence class: (1) separating threatened (CR, EN and VU) from non-threatened (NT and LC) species (binary classification); (2) separating CR species from other threatened species (EN and VU); (3) separating EN from VU in the remaining threatened species; and (4) separating NT from LC in the pool of non-threatened species. We repeated this modeling approach after excluding threatened species not categorized under criterion B (360 species), to explore the amount of uncertainty introduced by the other less commonly used Red List assessment criteria. Hyperparameter tuning and feature selection was performed at each classification task (description in Supporting Information).

**Figure 5.**
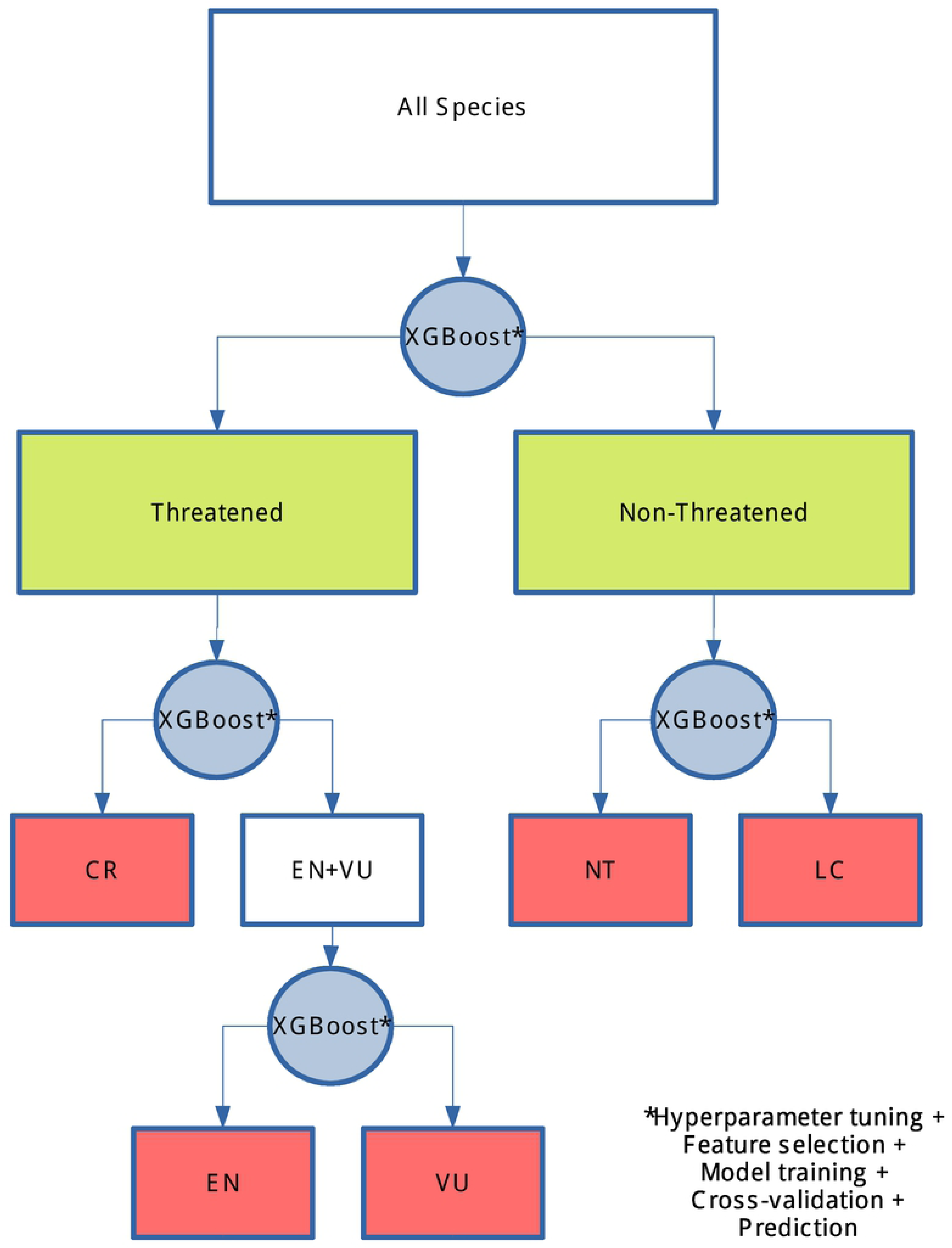
Flowchart for classification tasks in automated threat assessment method, using the XGBoost algorithm [27]. Green boxes represent outcomes of the binary task and red boxes represent the outcome of the specific tasks. Steps taken for each classification task (blue circle) are indicated after the asterisk.

Since supervised machine learning methods, such as XGBoost, are primarily predictive, rather than mechanistic, features contributing to better predictions are not necessarily useful for making causal inferences [28]. Thus, we evaluated the contribution of phylogenetic eigenvectors, Moran’s eigenvectors, and assessor/reviewer effects, by comparing models without these factors to models including them individually, and in different combinations (i.e. a model with only autocorrelations and a model with autocorrelations and assessor/reviewer effects-Table 1). This allowed us to explore if their inclusion increases predictive power. We also fit a model for the dataset excluding threatened species assessed by criteria other than B, but without assessor/reviewer effects as predictors, to evaluate the importance of these features on this subset of assessments. We plotted the number of previously evaluated species that changed from threatened to non-threatened categories and vice-versa, for each biogeographical realm [58], to evaluate spatial biases in the model errors.

### Comparison with previous methods

We also compared the features of our model to previously published automated assessment methods (incorporation of spatial and phylogenetic autocorrelation, assessor bias, tolerance to missing data, ability to predict specific IUCN categories). Beyond this we also implemented previous methods’ algorithms (where available), using our dataset of reptiles and predictors. These methods algorithms Random Forest [16,17], and Neural Networks [18,19], implemented using the R packages ’randomForest’ [59] and ’IUCNN’ [19], respectively. We compared these algorithms prediction accuracy with our method, in the binary task of separating threatened and non-threatened categories. We used the same predictors as in the XGBoost model, excluding the spatial and phylogenetic and assessor bias eigenvectors.

### Predictions for Data Deficient and Not Evaluated species

We used the model without assessor bias to estimate the threat categories of Data Deficient and Not Evaluated species. We used Pearson’s Chi-square to test if the proportions of DD and NE species predicted to be threatened were significantly different from the assessed ones. We further tested if proportions predicted for each threat category differ between DD, NE and assessed species. We adjusted p-values using False Discovery Rate [60].

### Phylogenetic and Spatial patterns

We explored how our predictions for DD and NE species changed the overall proportion of threatened species across the reptile phylogeny [30,31], different ecoregions [58], and biogeographical realms. For our phylogenetic representation we compared the proportion of threatened species in each clade before and after the addition our predictions for DD and NE species. We did this for all reptile families, as well as for each clade above the family level, and plotted the results along the branches of a composite phylogeny made from the trees of Tonini et al. [30] and Colston et al. [31].

We assigned species to ecoregions by intersecting species’ ranges from GARD 1.7 [41] with WWF terrestrial ecoregions of the world [58]. We compared the proportion of threatened species for each ecoregion, before and after the addition of predictions for DD and NE species. We also compared the percentage of threatened species before and after the inclusion of predictions for the eight terrestrial biogeographical realms: Afrotropics, Australasia, Indomalaya, Madagascar, Nearctic, Neotropics, Oceania, and Palearctic. Each species was assigned to all realms intersecting its range. The difference between proportions of threatened species in each biogeographical realm, before and after the inclusion of predictions, was tested using a Chi-square test, with p-values corrected for multiple comparisons, using False Discovery Rate [60].

### Effect of assessor/reviewer identities on predictions

We evaluated the effect of assessor/reviewer identities on predictions for each threat category. We sequentially permuted the assessor/reviewer eigenvector scores of each species to all other species, ran the modeling procedure described above, and retained the scores that resulted in least threatened (optimistic), and most threatened (pessimistic) categorizations. This procedure represents the potential results that would be obtained if the most “optimistic” and the most “pessimistic” group of assessors/reviewers assessed every species. This was done using the complete model using spatial and species-level predictors, spatial and phylogenetic autocorrelations, and assessor/reviewer effects, to minimize the effect of spatial and phylogenetic structure in assessor/reviewer assignments. We then tested if the resulting “optimistic” and “pessimistic” predictions were significantly different from the observed categories, and from each other, using Chi-square tests, with p-values corrected for multiple comparisons, using False Discovery Rate. We performed a similar analysis to explore differences in assessor effects with in each biogeographical realm for the binary classification task (of threatened/non-threatened categories).

## Acknowledgements

We thank all members of the Global Assessment of Reptile Distributions for making this work possible. The diligent and laborious work done by the IUCN global reptile assessment members. We thank Gopal Murali, Goni Barki, Anna Zimin, Anna Cihlová, Victor China, and Claudia Allegrini for fruitful discussions. We also thank two anonymous reviewers for valuable feedback that led to great improvement of this work.

## Supporting Information

**S1 Fig. Contribution of feature classes to the predictive performance of automated assessment models classifying reptile species into IUCN threat categories, for combinations of extent of occurrence class (columns, km^2^) and classification task (lines).** The ’Binary’ task separates threatened (CR, EN and VU) from non-threatened categories (NT and LC). CR – Critically Endangered, EN – Endangered, VU – Vulnerable, NT – Near Threatened, LC – Least Concern. Features in each class had their contribution measures summed. ’MEM’ stands for Moran’s Eigenvector Maps, an indicator of spatial autocorrelation. ’PEM’ stands for Phylogenetic Eigenvector Maps, an indicator of phylogenetic autocorrelation. For the specific identity of features in each class, see Supporting Information.

**S2 Fig. Number of reptile species in eight biogeographical realms that changed threat category after application of an automated assessment method, compared to the IUCN categories, under two categorization schemes:** a) binary (threatened vs non-threatened) categorization b) specific IUCN categories (CR, EN, VU, NT, LC). “Increases” indicates a species moved to a higher threat category, “decreases” indicates it moved to a lower threat category and “remains” indicates threat category stays the same. Y-axis is in log10 scale. AA – Australasian, AT – Afrotropical, IM – Indomalayan, MA – Madagascan, NA – Nearctic, NT – Neotropical, OC – Oceanian, PA – Palearctic.

**S3 Figure. Heatmap of threat category changes for different assessor bias scenarios.** Upper off diagonal elements represent the movements of species from less threatened to more threatened categories (left to right), in the pessimistic scenario. Lower off diagonal elements represent the movements of species from less threatened to more threatened categories (right to left), in the optimistic scenario. Diagonal indicates the IUCN threat categories species are moving to and from: CR – Critically Endangered, EN – Endangered, VU – Vulnerable, NT – Near Threatened, LC – Least Concern.

**S1 Table. Accuracy metrics of automated assessment models classifying reptile species into IUCN threat categories, under eight different approaches:** 1) complete model, accounting for spatial and phylogenetic autocorrelation and assessor/reviewer effects, 2) not accounting for spatial and phylogenetic autocorrelation or assessor/reviewer effects, 3) accounting for spatial autocorrelation, 4) accounting for phylogenetic autocorrelation, 5) accounting for spatial and phylogenetic autocorrelation, 6) accounting for assessor/reviewer effects, 7) accounting for spatial and phylogenetic autocorrelation and assessor/reviewer effects and excluding species categorized as threatened under criteria different from B, 8) accounting for spatial and phylogenetic autocorrelation and excluding species categorized as threatened under criteria different from B. ’Binary’ represents the separation of threatened (CR, EN and VU) from non-threatened categories (NT and LC). Remaining columns represent the predictive accuracy for assigning species to the five threat categories: CR – Critically Endangered, EN – Endangered, VU – Vulnerable, NT – Near Threatened, LC – Least Concern.

**S2 Table. Contribution of feature classes to the predictive performance of automated assessment models classifying reptile species into IUCN threat categories, for combinations of extent of occurrence class (km^2^) and classification task.** The ’Binary’ task separates threatened (CR, EN and VU) from non-threatened categories (NT and LC). CR – Critically Endangered, EN – Endangered, VU – Vulnerable, NT – Near Threatened, LC – Least Concern. Features in each class had their contribution measures summed. ’MEM’ stands for Moran’s Eigenvector Maps, an indicator of spatial autocorrelation. ’PEM’ stands for Phylogenetic Eigenvector Maps, an indicator of phylogentic autocorrelation. ’Assessors’ and ’reviewers’ stand for effects associated with the identity of assessors and reviewers that worked on each assessment. For the specific identity of features in each class, see Supporting Information.

**S3 Table. Optimal XGBoost hyperparameter configuration for each combination of classification tasks and extent of occurrence class.** Parameters adjusted were: learning rate (η), maximum tree depth (max_depth), minimum child weight (min_weight), row sampling (rowsample), column sampling (colsample), weight balancing (pos_weight), and three regularization parameters (γ, α, λ). Hyperparameter tuning strategy described in Supporting Information.

**S4 Table. Number of reptile species classified as threatened under non-B criteria in each IUCN category before (rows) and after (columns) application of automated assessment method trained on B criteria species.**

**S5 Table. Accuracy metrics of two previously published automated assessment models for separating reptile species into threatened (CR, EN and VU) and non-threatened categories (NT and LC) IUCN threat categories.** Random Forest refers to the approach described by Bland et al, 2015, and Neural networks refers to the approach described by Zizka et al, 2020.

**S6 Table. Comparison of automated assessment methods.** Models are compared in their incorporation of spatial and phylogenetic autocorrelation, as well as their ability to account for assessor bias, including missing data and predicting specific IUCN categories. The method presented here is indicated as Caetano et al., 2021.

**S7 Table. Pearson’s Χ^2^ test statistics for comparisons of the proportion of reptile species assigned to each IUCN category between the actual assessments (Observed) and the predictions for Data Deficient (DD) and Not Evaluated species (NE) species, made using an automated assessment model.** We adjusted p-values adjusted for False Discovery Rate. ’Threatened’ represents the proportion of species assigned a threatened category (CR, EN and VU). CR – Critically Endangered, EN – Endangered, VU – Vulnerable, NT – Near Threatened, LC – Least Concern. Significant p-values are in bold.

**S8 Table. Number of reptile species in each IUCN category before (rows) and after (columns) application of automated assessment method.**

**S9 Table. Difference in the proportion of threatened species in reptile families before and after the addition of threat estimates for Data Deficient and Not Evaluated species, obtained from an automated assessment method.**

**S10 Table. Pearson’s Χ^2^ test statistics for comparisons of the proportion of threatened reptile species in eight biogeographical realms, before and after the inclusion of predictions for Data Deficient and Not Evaluated species, made using an automated assessment model.** We adjusted p-values adjusted for False Discovery Rate. Significant p-values are in bold.

**S11 Table. Pearson’s Χ^2^ test statistics for comparisons of the proportion of reptile species assigned to each IUCN category between the actual assessments (Observed) and the expected if the most optimist group of assessors assessed every species (Optimist) and if the most group pessimist assessed every species (Pessimist), estimated using an automated assessment model.** We adjusted p-values adjusted for False Discovery Rate. ’Threatened’ represents the proportion of species assigned a threatened category (CR, EN and VU). CR – Critically Endangered, EN – Endangered, VU – Vulnerable, NT – Near Threatened, LC – Least Concern. Significant p-values are in bold.

**S12 Table. Pearson’s Χ^2^ test statistics for comparisons of the proportion of threatened reptile species in eight biogeographical realms between the actual assessments (Observed) and the expected if the most optimist group of assessors assessed every species (Optimist) and if the most group pessimist assessed every species (Pessimist), estimated using an automated assessment model.** We adjusted p-values adjusted for False Discovery Rate.

